## Supplementary Information for "Structure of full-length cobalamin-dependent methionine synthase and cofactor loading captured *in crystallo*"

#### Supplementary Figures

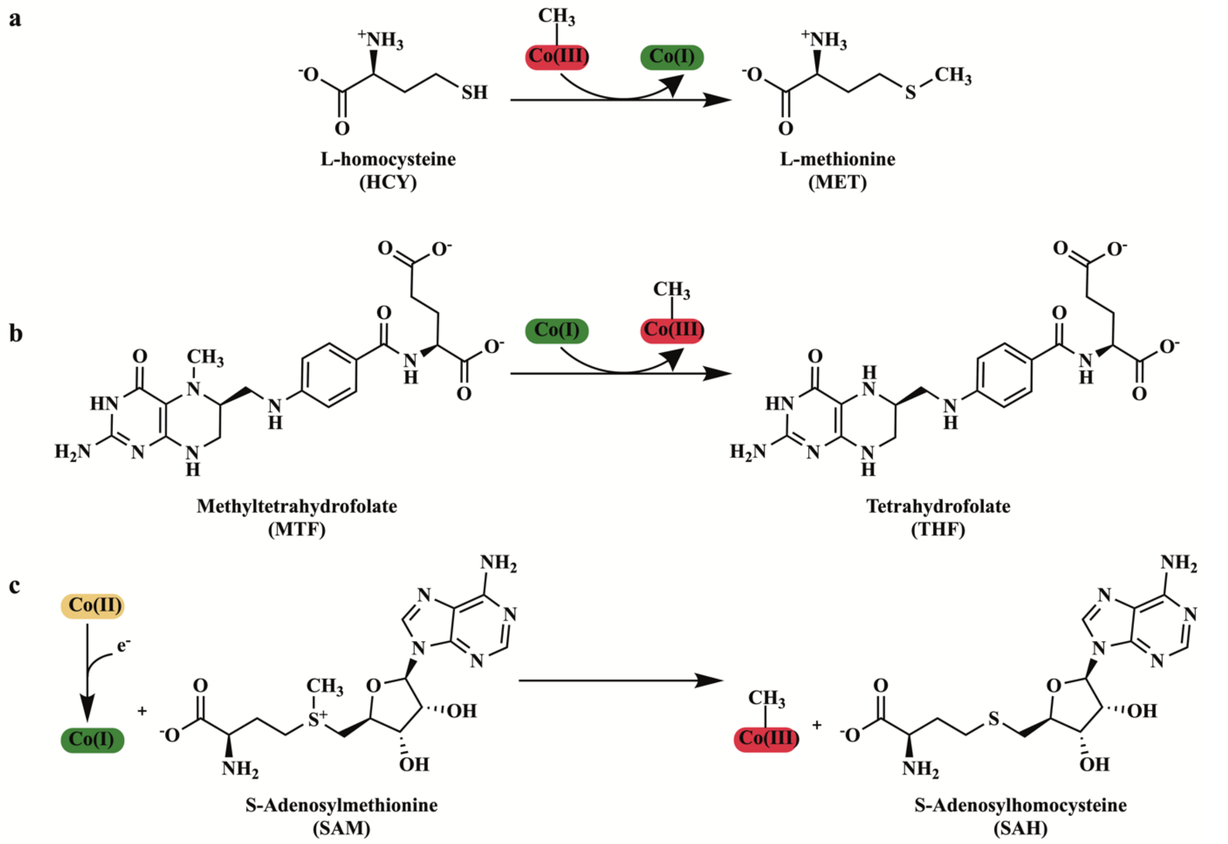


**Supplementary Figure 1. The three methylations catalyzed by methionine synthase.** **a** The methylation of homocysteine by MS using $CH_3$-Co(III) (MeCbl) to form methionine. **b** The demethylation of methyltetrahydrofolate by MS using Co(I) to yield $CH_3$-Co(III) (MeCbl). **c** The reactivation reaction catalyzed by MS, a reductive methylation and reactivation of Co(II) to yield $CH_3$-Co(III) (MeCbl).

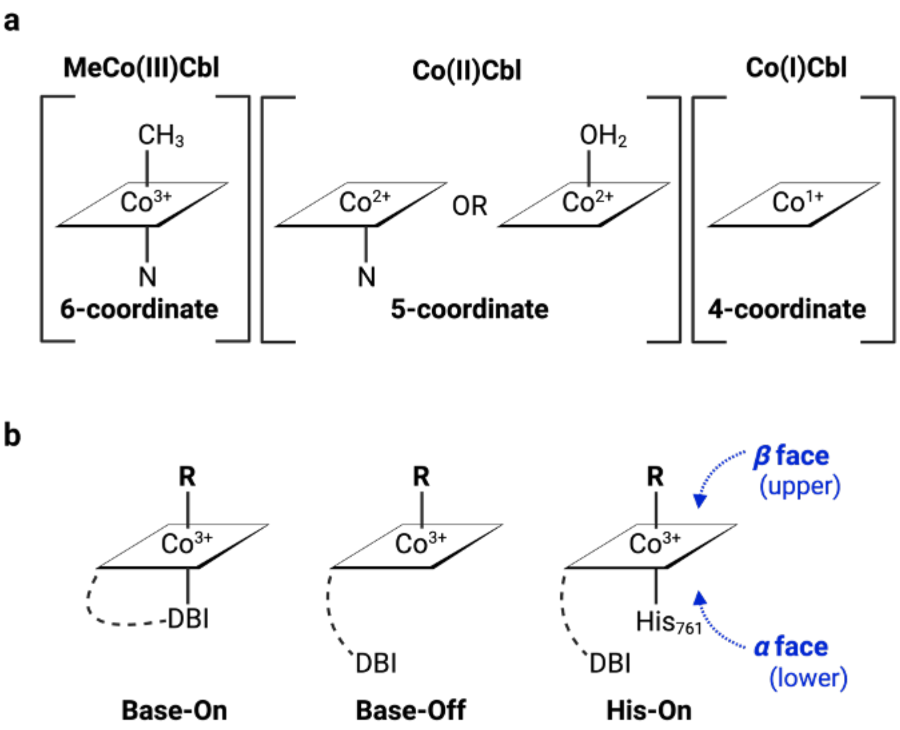


**Supplementary Figure 2. Cobalamin coordination environment. a** Coordination environment of cobalamin in different cobalt oxidation states. **b** Base-on/Base-off. In solution, the DBI tail is coordinated in the lower axial position. Upon binding, the DBI tail is replaced with His to achieve the His-on state.

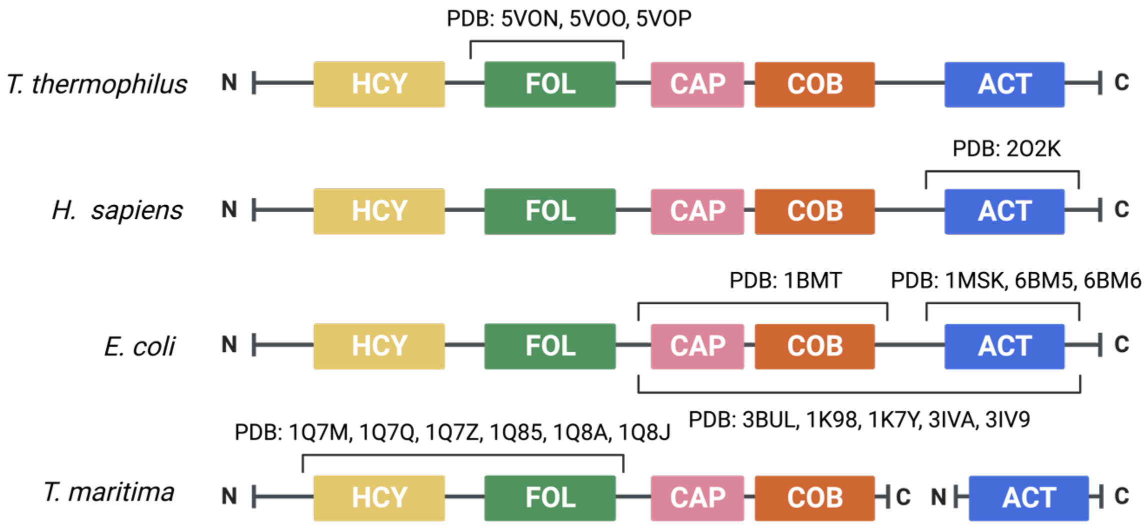


**Supplementary Figure 3. MS domain organization and PDB structures.**

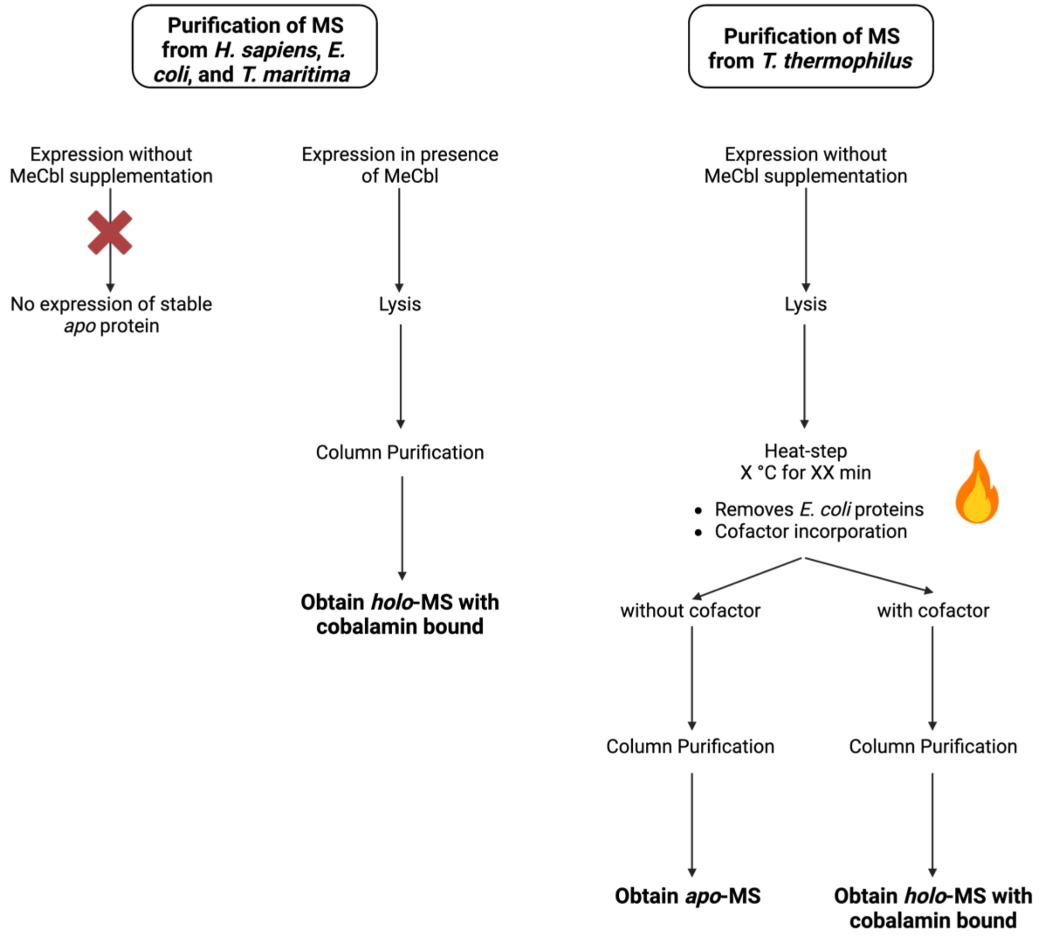


**Supplementary Figure 4. MS versus *t*MS purification workflow.**

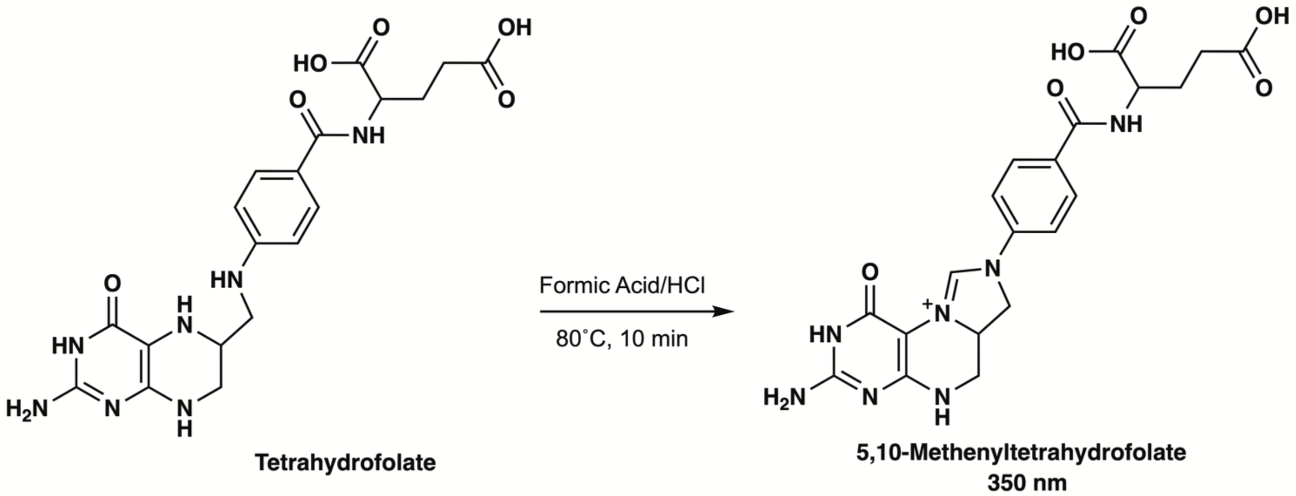


**Supplementary Figure 5. Tetrahydrofolate acid-catalyzed conversion to methenyltetrahydrofolate.** The resulting product has a characteristic absorbance at 350 nm and is used to track *t*MS activity in a coupled-assay described in Methods.

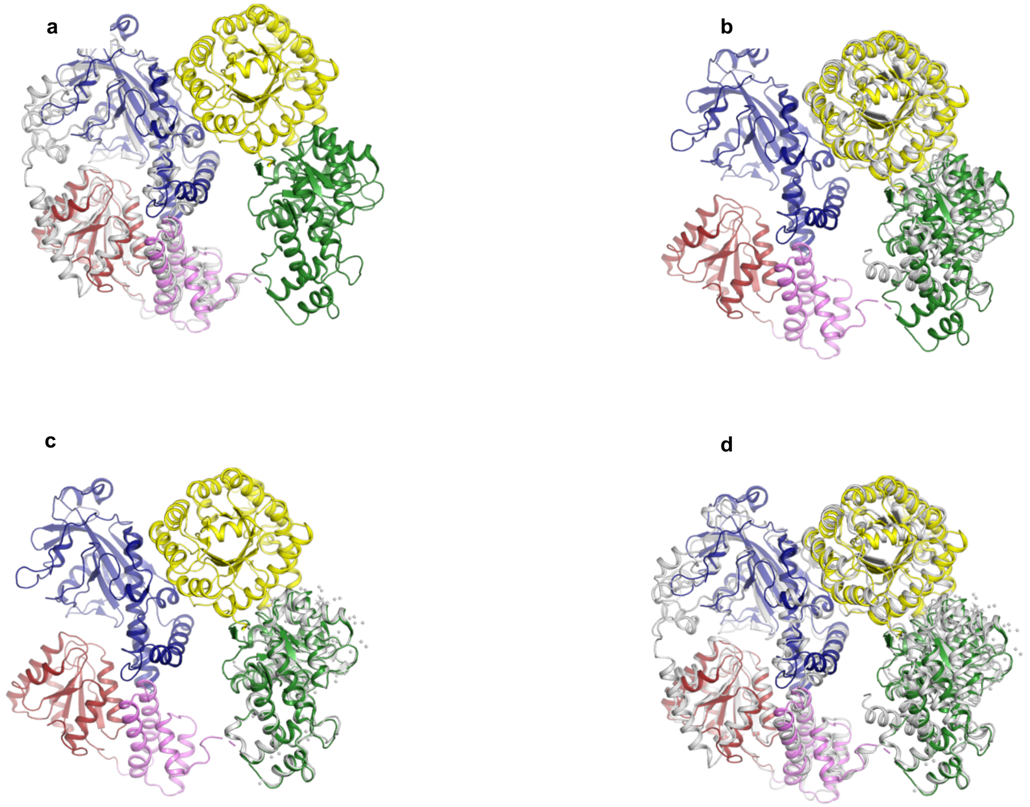


**Supplementary Figure 6. Alignment excised MS domains with full-length *t*MS. a** Alignment of Cap:Cob:Act domains from 1K7Y (gray) with Cap (pink), Cob (red), and Act (blue) in full-length *t*MS – RMSD=2.16 Å. **b** Alignment of Fol and Hcy domains from 3BOL (gray) with the Fol (green) and Hcy (yellow) domains in full-length *t*MS – RMSD=2.64 Å. **c** Alignment of Fol 5VON (gray) domain with Fol domain (green) in full-length *t*MS – RMSD=0.46 Å. **d** Alignment of the excised domains (gray) from 1K7Y, 3BOL, and 5VON with full-length *t*MS.

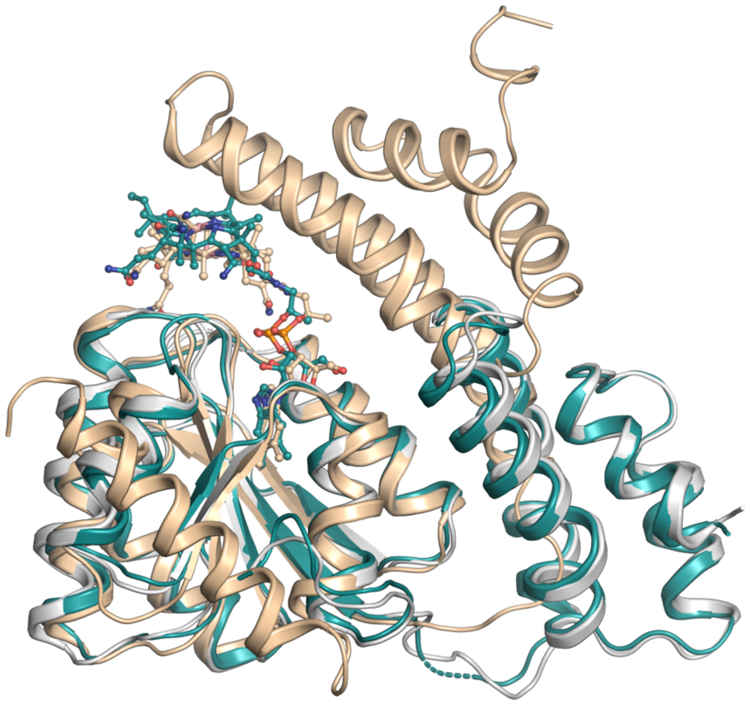

**Supplementary Figure 7**. **Displacement of Cap-domain in full length MS.** Alignment of Cap:Cob domains from the full length structure (gray, reactivation) with the Cob domain from *holo*-Cap:Cob:Act (teal, reactivation, His-off) – RMSD=0.72 Å and the Cap:Cob domains from 1BMT (wheat, reactivation, His-on) – RMSD=0.85 Å.

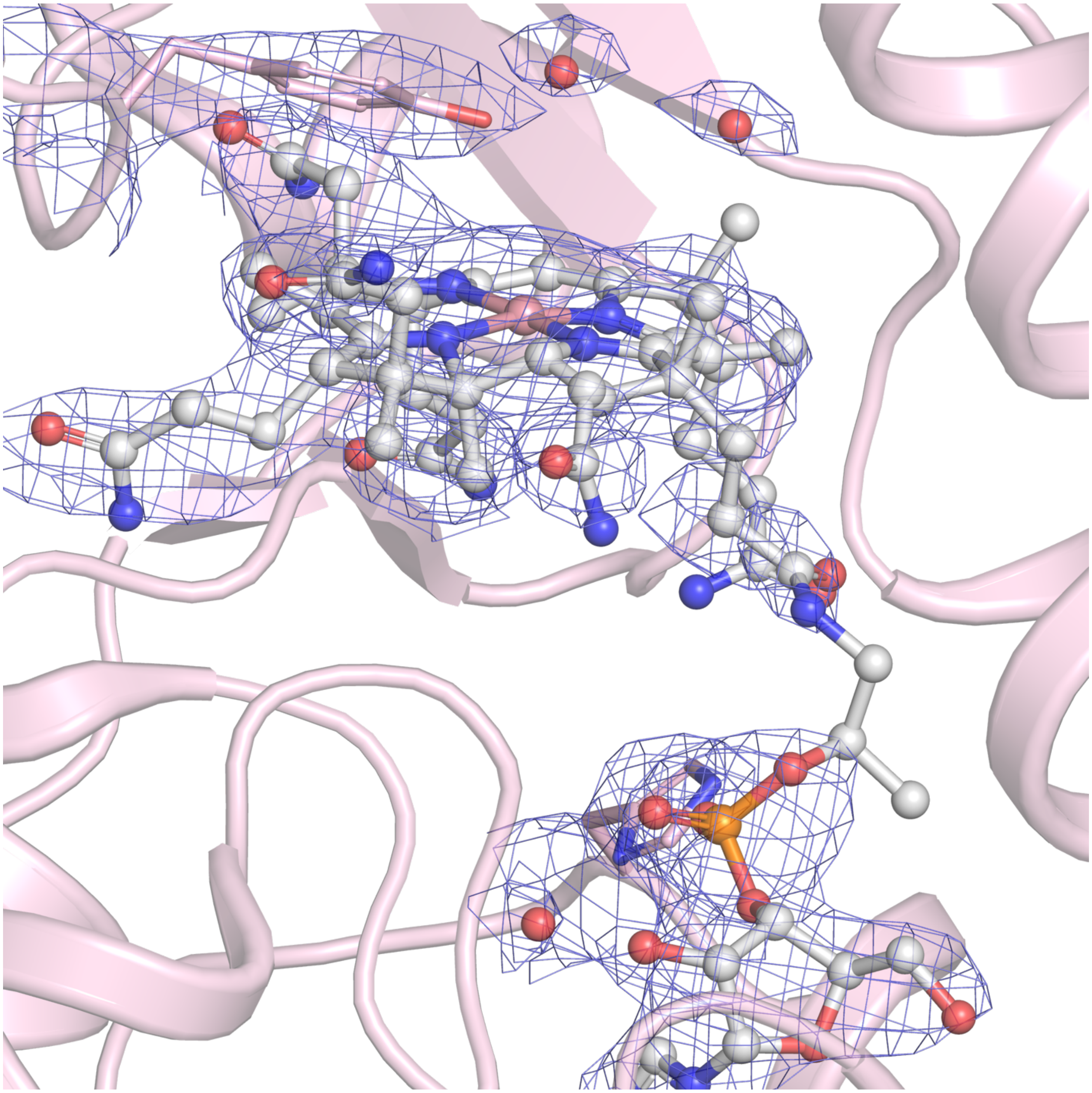

**Supplementary Figure 8. *t*MS electron density around the cobalamin cofactor.** *Holo*-Cap:Cob:Act (light pink) and its cobalamin cofactor (gray), along with Tyr1132 and His761 in the axial cofactor positions. Their corresponding electron density ($2F_o$-$F_c$) contoured at 1.5 σ are shown in blue.

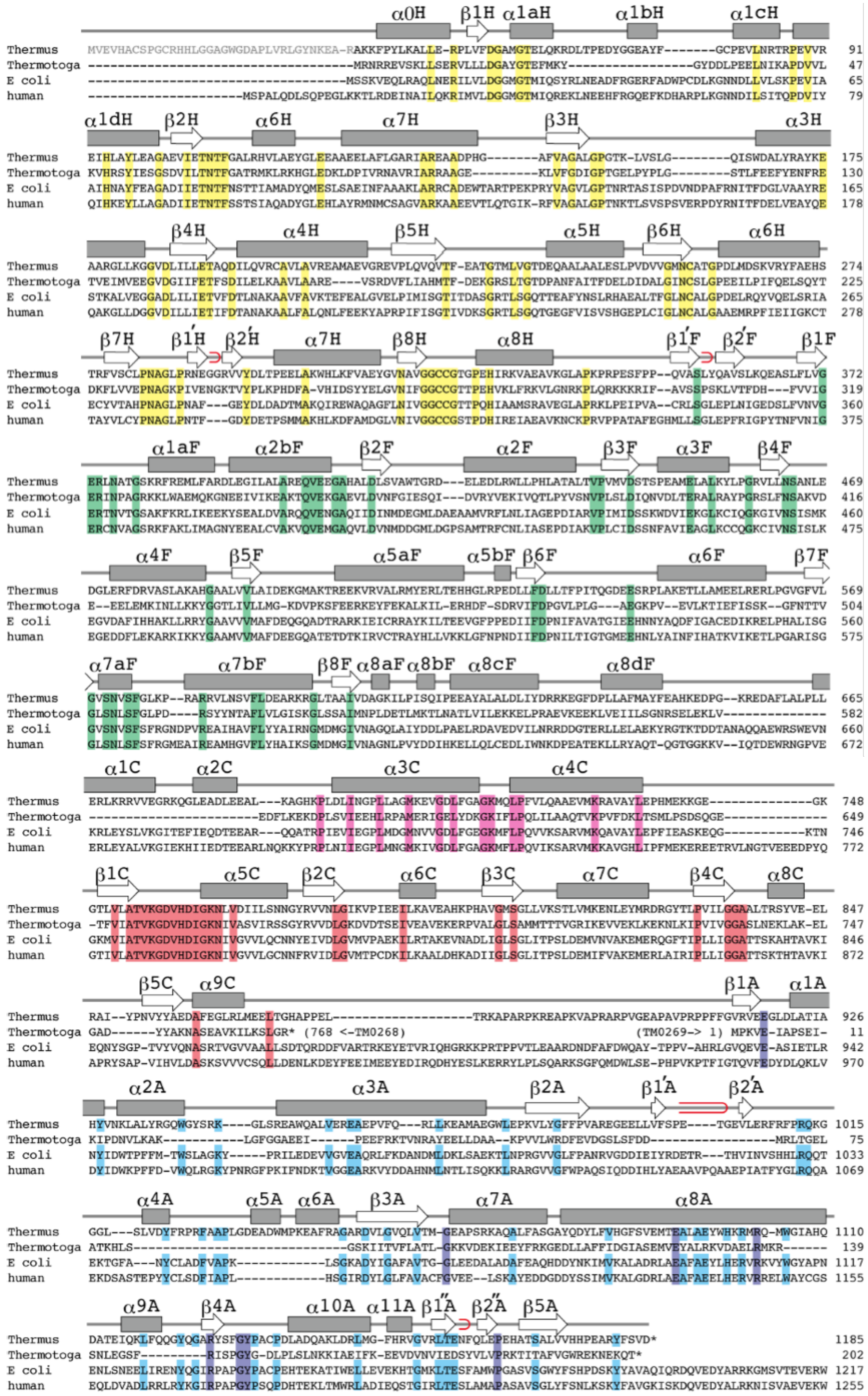

α0H β1H α1aH α1bH α1cH
α1dH β2H α6H α7H β3H α3H
β4H α4H β5H α5H β6H α6H
β7H β1'H β2'H α7H β8H α8H β1'F β2'F β1F
α1aF α2bF β2F α2F β3F α3F β4F
α4F β5F α5aF α5bF β6F α6F β7F
α7aF α7bF β8F α8aF α8bF α8cF α8dF
α1C α2C α3C α4C
β1C α5C β2C α6C β3C α7C β4C α8C
β5C α9C β1A α1A
α2A α3A β2A β1'A β2'A
α4A α5A α6A β3A α7A α8A
α9A β4A α10A α11A β1''A β2''A β5A
Thermus
Thermotoga
E coli
human
(768 <-TM0268)
(TM0269-> 1)

**Supplementary Figure 9. Methionine synthase (MS) sequence alignment.** Schematic illustration of the secondary structure of *t*MS and alignments with the amino acid sequence of MS from *Thermus thermophilus* [Genbank accession number NC_00646], *Thermotoga maritima* [NC_000853], *Escherichia coli* [J04975], and human [U73338]. α-helices and β-sheets are shown in boxes and arrows, respectively. Red loops indicate β-hairpins. The N-terminal residues absent in *t*MS$^{\Delta N35}$ are shown in gray. Conserved amino acid residues are highlighted by yellow, green, pink, red, and blue for the homocysteine, folate, Cap, cobalamin, and *S*-adenosylmethionine domains, respectively. In the activation (Act) domain, cyan is used to highlight conserved amino acid residues in *T. thermophilus*, *E. coli*, and human MS.

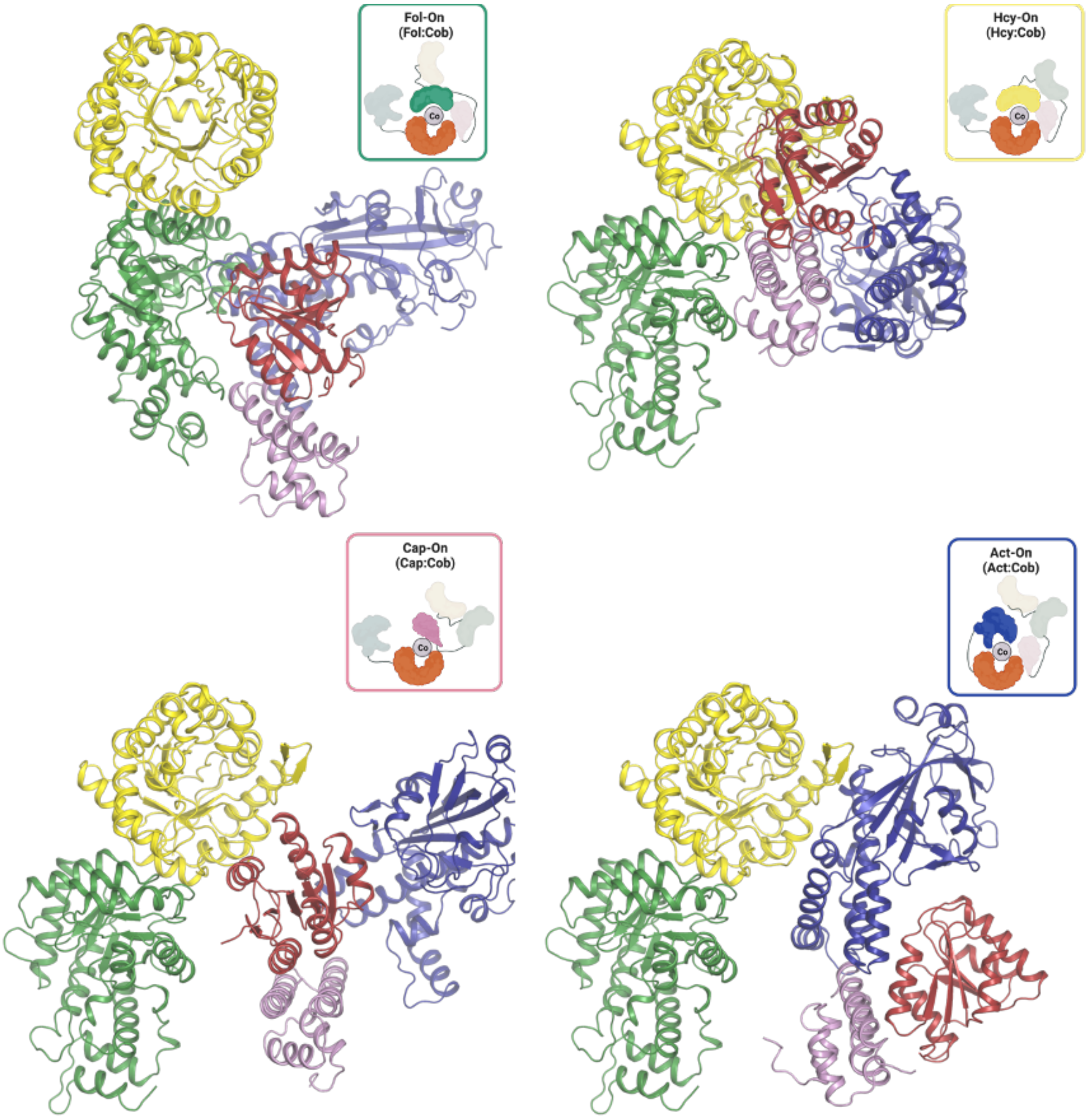


**Supplementary Figure 10. Conformational dynamics of MS.** The proposed conformations adopted by MS throughout the catalytic and reactivation cycles with different substrate binding domains positioned above the cofactor; Fol-on (top-left), Hcy-on (top-right), Cap-on (bottom-left), and Act-on (bottom-right).

#### Supplementary Tables

##### Supplementary Table 1. Kinetic parameters of Methionine Synthase

| Enzyme | Molecular Weight (kDa) | Specific Activity (μmol $min^{-1}$ $mg^{-1}$ protein) | $k_{cat}$ ($min^{-1}$) | $K_m$ (μM) | |
|---|---|---|---|---|---|
| | | | | (6S)$CH_3$-$H_4$folate | Homocysteine |
| *E. coli* MS | 133 | 11.6[1] | 1542 at 37°C<br>1128 at 25°C[2] | 27.8 | 0.8 |
| *T. thermophilus* MS | 131 | 8.1 | 1062 at 50°C | 18.4 ± 4.1 | 9.3 ± 2.8 |
| *S. scrofa* MS (Pig) | 150 | 1.7 | 255 | 12.6 | 2.16 |
| *R. rattus* MS (Rat) | 140 | 1.6 | 224 | 38 | 1.7 |

[1]Biochemistry 1998, 27, 8458-65
[2]Biochemistry 1990, 29, 11101-9

**Supplementary Table 2. X-Ray Data Collection and Refinement Statistics for *t*MS$^{\Delta N35}$, *Apo-t*MS$^{Cap:Cob:Act}$, and *Holo-t*MS$^{Cap:Cob:Act}$**

| | ***t*MS$^{\Delta N35}$** | ***Apo-t*MS$^{Cap:Cob:Act}$** | ***Holo-t*MS$^{Cap:Cob:Act}$** |
|---|---|---|---|
| **Data collection** | | | |
| Beamline | APS, GMCA 23-IDB | APS, GMCA 23-IDB | APS, GMCA 23-IDB |
| Wavelength (Å) | 1.033 | 1.033 | 1.033 |
| Temperature (K) | 100 | 100 | 100 |
| Resolution (Å) | 106.64-2.75 (2.85-2.75) | 40.55-2.40 (2.50-2.40) | 49.30-3.15 (3.15-3.23) |
| Space group | $P4_12_12$ | $P3_121$ | $C121$ |
| Cell dimensions (Å) | a = b = 134.84, c = 174.74 | a = b = 96.18, c = 356.04 | a = 166.135, b = 95.844, c = 238.745 |
| Cell dimensions (°) | α = β = γ = 90 | α = γ = 90, β = 120 | α = γ = 90, β = 91.96 |
| Observed reflections | 669,916 (51,333) | 711,608 (42,258) | 222,439 (11,567) |
| Unique reflections | 42,467 (4,373) | 76,051 (4,396) | 64,264 (3,294) |
| $R_{meas}$ (%) | 19.7 (167.3) | 18.7 (95.4) | 22.0 (116.7) |
| $R_{merge}$ (%) | 19.1 (160.0) | 17.7 (90.3) | 18.5 (98.6) |
| $\langle I/\sigma \rangle$ | 12.3 (1.8) | 8.1 (2.5) | 5.1 (1.3) |
| CC(1/2) | 0.993 (0.672) | 0.99 (0.667) | 0.99 (0.581) |
| Multiplicity | 15.8 (11.7) | 9.4 (9.6) | 3.5 (3.5) |
| Completeness (%) | 100.00 (100.00) | 100.00 (100.00) | 98.50 (99.22) |
| Overall *B* (Å$^2$) (Wilson plot) | 56.2 | 47.00 | 60.70 |
| **Refinement** | | | |
| Resolution range | 106.96 - 2.75 | 39.85 - 2.95 | 49.30 - 3.15 |
| Number of reflections (work/test set) | 40,263/2,154 | 75,954/3,830 | 64,264/3,294 |
| $R_{work}/R_{free}$ (%) | 22.6/25.3 | 26.5/29.9 | 24.6/30.1 |
| No. of non-H atoms | | | |
| Protein | 8751 | 12,223 | 24,885 |
| Water | 131 | 493 | 299 |
| Ligand | 11 | 0 | 546 |
| B-factors (Å$^2$) | | | |
| Protein | 81.63 | 40.63 | 84.67 |
| Water | 58.80 | 44.00 | 44.50 |
| Ligand | 133.13 | 0.00 | 70.93 |
| Rmsd deviations | | | |
| Bond lengths (Å) | 0.008 | 0.0075 | 0.012 |
| Bond angles (°) | 1.36 | 1.70 | 1.99 |
| Favored/allowed/outliers | 96.9/3.0/0.1 | 97.5/2.5/0.0 | 97.9/2.1/0.0 |
| MolProbity Score | 1.03 (100$^{th}$ percentile) | 0.60 (100$^{th}$ percentile) | 0.99 (100$^{th}$ percentile) |
| PDB | 8SSC | 8SSD | 8SSE |

**Supplementary Table 3. Purification Yield and Activity**

| | Protein* (mg) | Total activity (µmol $min^{-1}$) | Specific Activity (µmol $min^{-1}$ $mg^{-1}$ protein) | Purification *n*-Fold | Yield (%) |
|---|---|---|---|---|---|
| Crude Extract | 674# | 700 | 1.04 | 1.0 | 100 |
| Heat treatment | 218 | 390 | 1.80 | 1.7 | 56 |
| Ni-affinity Chromatography | 37 | 300 | 8.11 | 8.0 | 44 |

*Protein concentration was determined using the Bradford method {Bradford, **1976**, 2300}.
### $CH_3$-cobalamin was added when activity was measured.
